# 3D Perception of Maximum Density Zone on Ramachandran Plots for Zika Virus Protein Structures

**DOI:** 10.1101/053074

**Authors:** Mayukh Mukhopadhyay, Parama Bhaumik

## Abstract

The Ramachandran plot is among the most central concepts in structural biology which uses torsion angles to describe polypeptide and protein conformation. To help visualize the features of high-fidelity Ramachandran plots, it is helpful to look beyond the common two-dimensional psi-phi-plot, which for a large dataset does not serve very well to convey the true nature of the distribution. In particular, when a large subset of the observations is found very narrowly distributed within one small region, this is not well seen in the simple plot because the data points congest one another. Zika Virus (ZIKV) protein databank has been chosen as specimen for analysis. This is because the structure, tropism, and pathogenesis of ZIKV are largely unknown and are the focus of current investigations in an effort to address the need for rapid development of vaccines and therapeutics. After a brief survey on Zika Virus, it is shown that when a dense dataset of ZIKV protein databank is passed through a colour-coded scaled algorithm, a three dimensional plot gets generated which gives a much more compelling impression of the proportions of residues in the different parts of the protein rather than representing it in a normal two dimensional psi-phi plot.

## I. Brief Survey On Zika Virus

### 1.1. Introduction

In May 2015 an outbreak of Zika virus began in Brazil and spread across many regions of Central and South America. The spread of the virus led to reports of health problems such as Guillain-Barré syndrome and possibly microcephaly, a disease in which babies are born with abnormally small heads and brains that have not developed properly [10]. The WHO said a direct causal relationship between Zika virus infection and birth defects has not yet been established but is strongly suspected. However, Dr. Tom Frieden, director of the U.S. Centers for Disease Control and Prevention (CDC) reports that “There isn't any doubt that Zika causes microcephaly." As a result, the CDC issued travel advisories for those traveling to affected areas [9].

### 1.2. Description and Significance

Zika virus (ZIKV) is a mosquito-borne flavivirus that was first isolated from a rhesus monkey in the Zika forest of Uganda in 1947 [5]. In 1968, isolation from human hosts occurred in residents of Nigeria [6]. Since then, multiple studies have confirmed ZIKV antibody in humans from a multitude of countries in Africa and parts of Asia [6]. In 2015, ZIKV first appeared outside of Africa and Asia when it was isolated in Brazil where is has caused a minor outbreak following the 2014 FIFA World Cup [7]. ZIKV is closely related to other mosquito-borne flaviviruses such as the dengue, yellow fever, West Nile, and Japanese encephalitis viruses [6]. ZIKV causes a disease known as Zika fever, which is characterized by a macropapular rash covering the body, fever, joint pain, and malaise [6]. Although there have yet to be serious complications arising from ZIKV, its appearance across the globe, mosquito-driven transmission cycle, and possible spread via sexual contact make ZIKV an important emerging pathogen whose global impact is yet to be discovered [11]. Figure 1 illustrates the spread of Zika virus.

**Figure 1.**
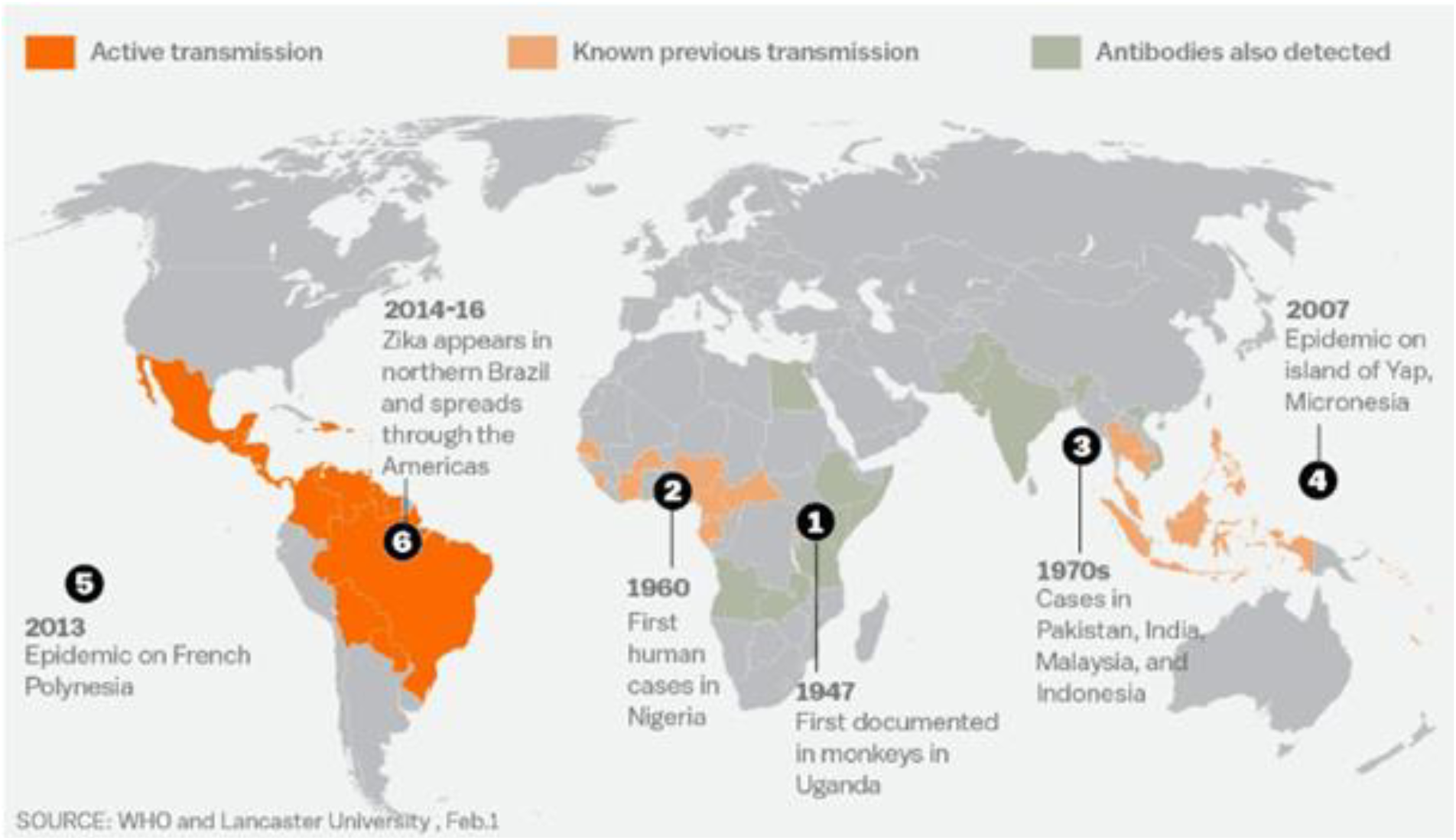
Spread of ZIKV [11]

### 1.3. Genome Structure

The Zika virus is a positive sense single-stranded RNA molecule 10794 bases long with two non-coding regions flanking regions known as the 5' NCR and the 3' NCR [5]. The open reading frame of the Zika virus reads as follows: 5′-C-prM-E-NS1-NS2A-NS2B-NS3-NS4A-NS4B-NS5-3′ and codes for a polyprotein that is subsequently cleaved into capsid (C), precursor membrane (prM), envelope (E), and non-structural proteins (NS) [8]. The E protein composes the majority of the virion surface and is involved with aspects of replication such as host cell binding and membrane fusion. NS1, NS3, and NS5 are large, highly-conserved proteins while the NS2A, NS2B, NS4A, and NS4B proteins are smaller, hydrophobic proteins. Located in the 3' NCR are 428 nucleotides that may play a part in translation, RNA packaging, cyclization, genome stabilization, and recognition. The 3' NCR forms a loop structure and the 5' NCR allows translation via a methylated nucleotide cap or a genome-linked protein [12].

### 1.3. Virion Structure of Zika Virus

The structure of ZIKV follows that of other flaviviruses. It contains a nucleocapsid approximately 25-30 nm in diameter surrounded by a host-membrane derived lipid bilayer that contains envelope proteins E and M. The virion is approximately 40 nm in diameter with surface projections that measure roughly 5-10 nm [8]. The surface proteins are arranged in an icosohedral-like symmetry [12].

As part of our analysis we have chosen the following specimens currently available in RCSB protein databank:

a. The cryo-EM structure of Zika Virus (PDB ID 5IRE)
b. ZIKV nonstructural protein 1 (NS1) (PDB ID 5IY3)

#### 1.3.1 The cryo-EM structure of Zika Virus (PDB ID 5IRE)

A cryo–electron microscopy (cryo-EM) structure of the mature ZIKV at near-atomic resolution (3.8 Å) has been reported in the protein data bank [3]. The structure of Zika virus is similar to other known flavivirus structures, except for the ~10 amino acids that surround the Asn154 glycosylation site in each of the 180 envelope glycoproteins that make up the icosahedral shell [3].

The carbohydrate moiety associated with this 5IRE residue, which is recognizable in the cryo-EM electron density, may function as an attachment site of the virus to host cells [3].

Figure 4 summarizes the cryo-EM structure of ZIKV at 3.8 Å. (A) A representative cryo-EM image of frozen hydrated ZIKV, showing the distribution of virion phenotypes. Smooth, mature virus particles are shown enclosed in black boxes. A partially mature virus particle is identified by the yellow arrow. (B) A surface-shaded depth-cued representation of ZIKV, viewed down the icosahedral twofold axis. The asymmetric unit is identified by the black triangle. (C) A cross section of ZIKV showing the radial density distribution. Colour coding in (B) and (C) is based on radii, as follows: blue, up to 130 Å; cyan, 131 to 150 Å; green, 151 to 190 Å; yellow, 191 to 230 Å; red, 231 Å and above. The region shown in blue fails to follow icosahedral symmetry, and therefore its density is uninterruptable, as is the case with other flaviviruses. (D) A plot of the Fourier shell correlation (FSC). Based on the 0.143 criterion for the comparison of two independent data sets, the resolution of the reconstruction is 3.8 Å. (E) The Cα backbone of the E and M proteins in the icosahedral ZIKV particle [same orientation as in (B)], showing the herringbone organization. The colour code follows the standard designation of E protein domains I (red), II (yellow), and III (blue). (F) Representative cryo-EM electron densities of several amino acids of the E protein. Cyan indicates carbon atoms; dark blue, nitrogen atoms; red, oxygen atoms; and yellow, sulfur atoms. Single-letter abbreviations for the amino acid residues are as follows: A, Ala; C, Cys; D, Asp; E, Glu; F, Phe; G, Gly; H, His; I, Ile; K, Lys; L, Leu; M, Met; N, Asn; P, Pro; Q, Gln; R, Arg; S, Ser; T, Thr; V, Val; W, Trp; and Y, Tyr.

**Figure 2.**
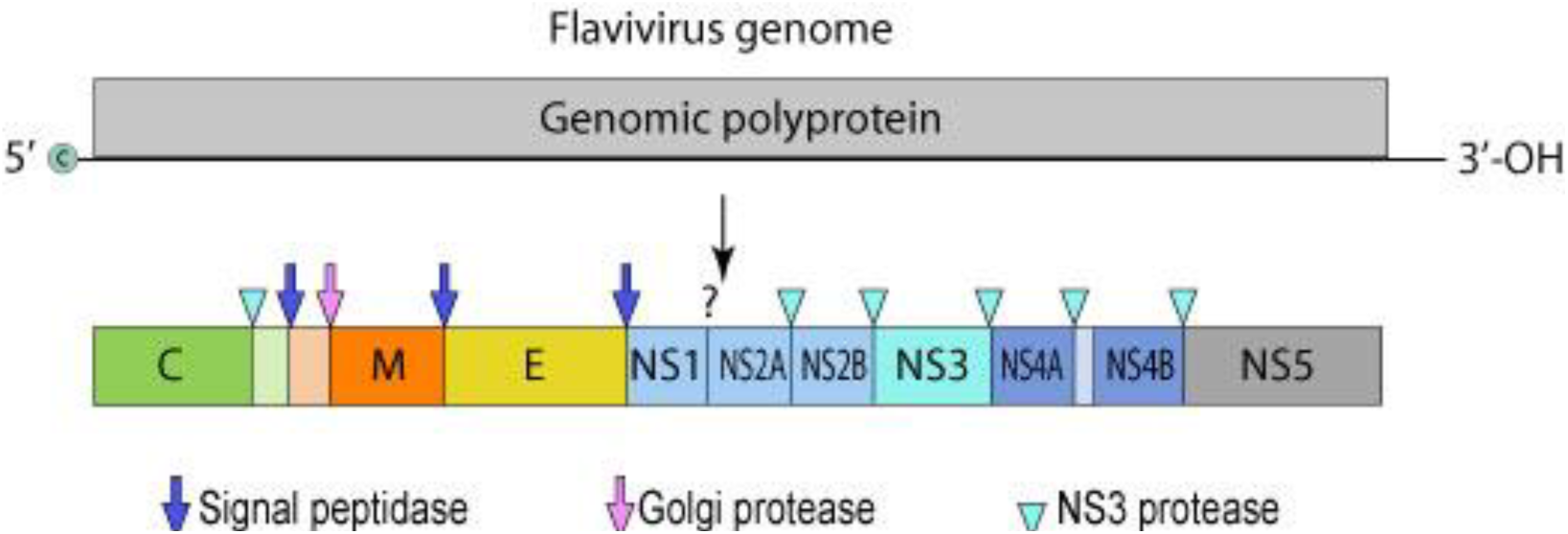
General Genome Structure of Flavivirus [7]

**Figu0re 3.**
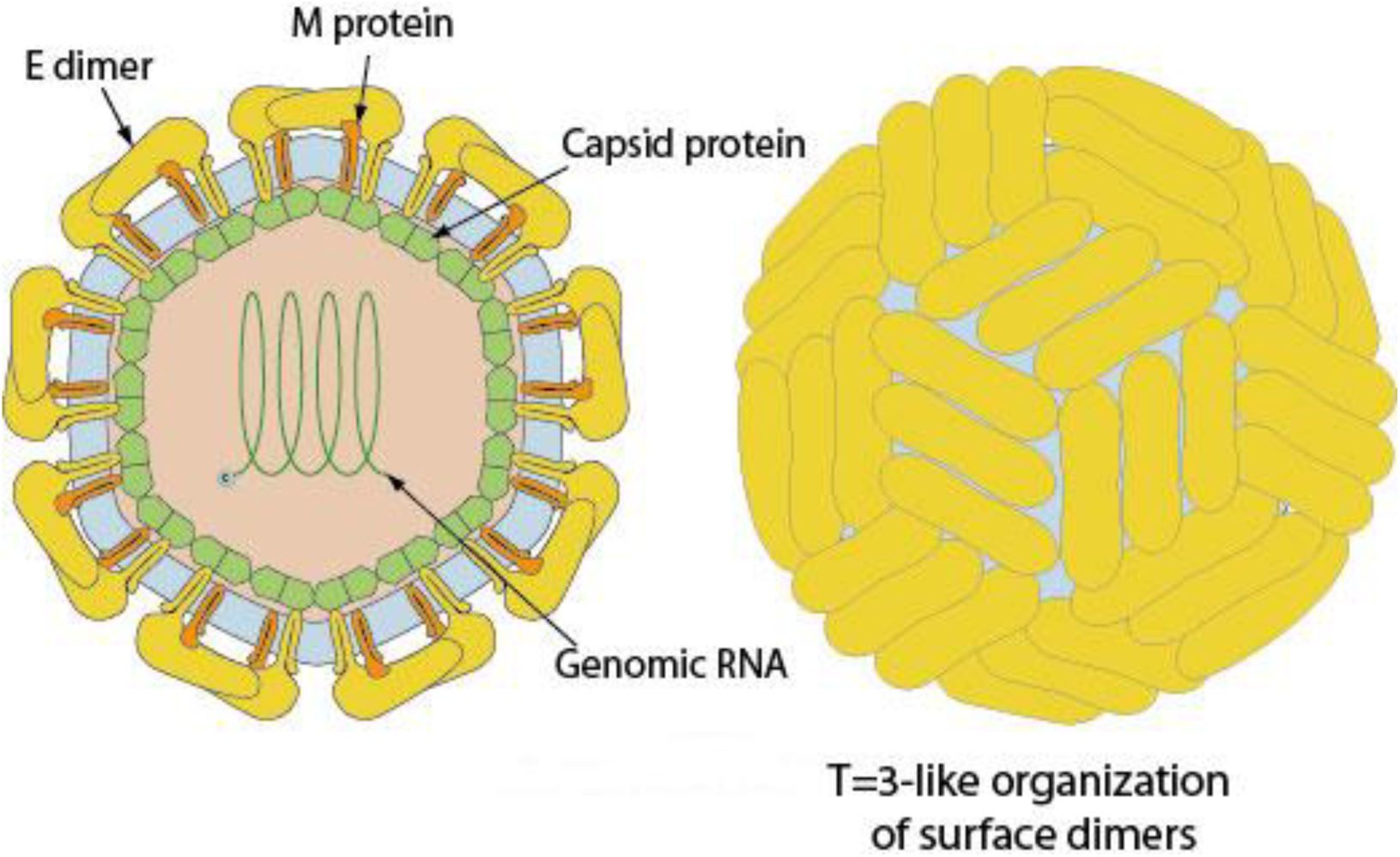
Virion structure of Flavivirus [7]

**Figure 4.**
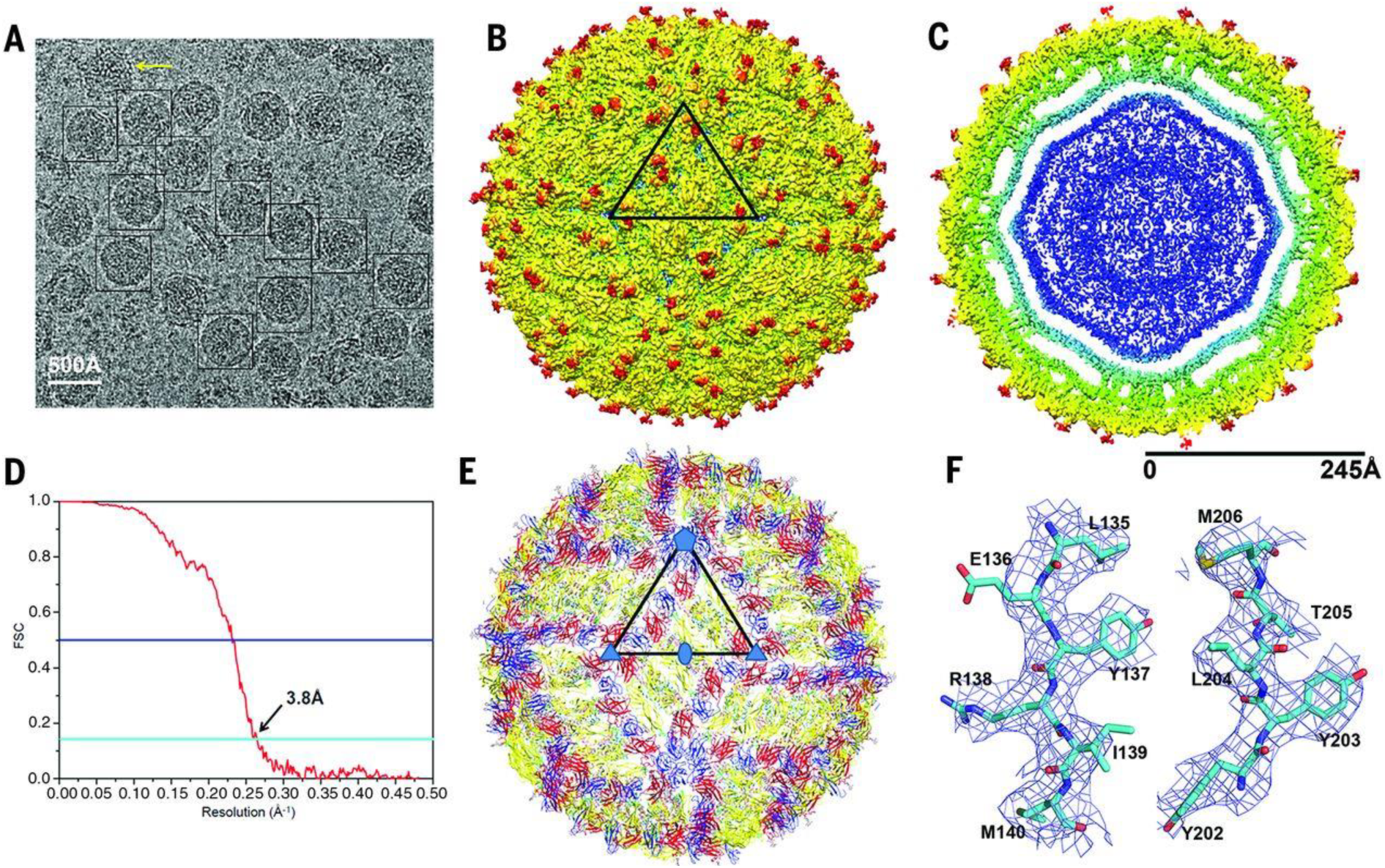
3.8 Å resolution cryo-EM structure of Zika virus [3]

#### 1.3.2 ZIKV non-structural protein 1 (NS1) (PDB ID 5IY3)

The molecular mechanisms of NS1 are relatively well established for Dengue Virus (DENV) and West Nile virus (WNV) and the NS1-encoding sequence is suspected to be a major genetic factor underlying the diverse clinical consequences of infections caused by flaviviruses (over 70 members). However, little is known about the NS1 of ZIKV, which displays different pathogenesis from that of typical flaviviruses [2].

In the protein data bank, virologists have address this lack of information, by expressing the ZIKV NS1172–352 fragment of the BeH819015 strain isolated from Brazil in 2015 in Escherichia coli as inclusion bodies and obtained the soluble protein by in vitro refolding and then solved its crystal structure by molecular replacement to a resolution of 2.2 Å [2].

The ZIKV NS1172–352 protein crystallized as a rod-like homodimer with a length of ~9 nm. Sedimentation velocity analytical ultracentrifugation analyses confirmed that the ZIKV NS1172–352 protein exists as a homodimer (~40 kDa) in solution [2].

In the below figure, The ZIKV NS1172–352 homodimer structure has a continuous β-sheet on one surface, with 20 β-strands arranged like the rungs of a ladder, in which each monomer contributes ten rungs to the antiparallel β-ladder. On the opposite side of the homodimer, an irregular surface is formed by a complex arrangement of loop structures [2].

**Figure 5.**
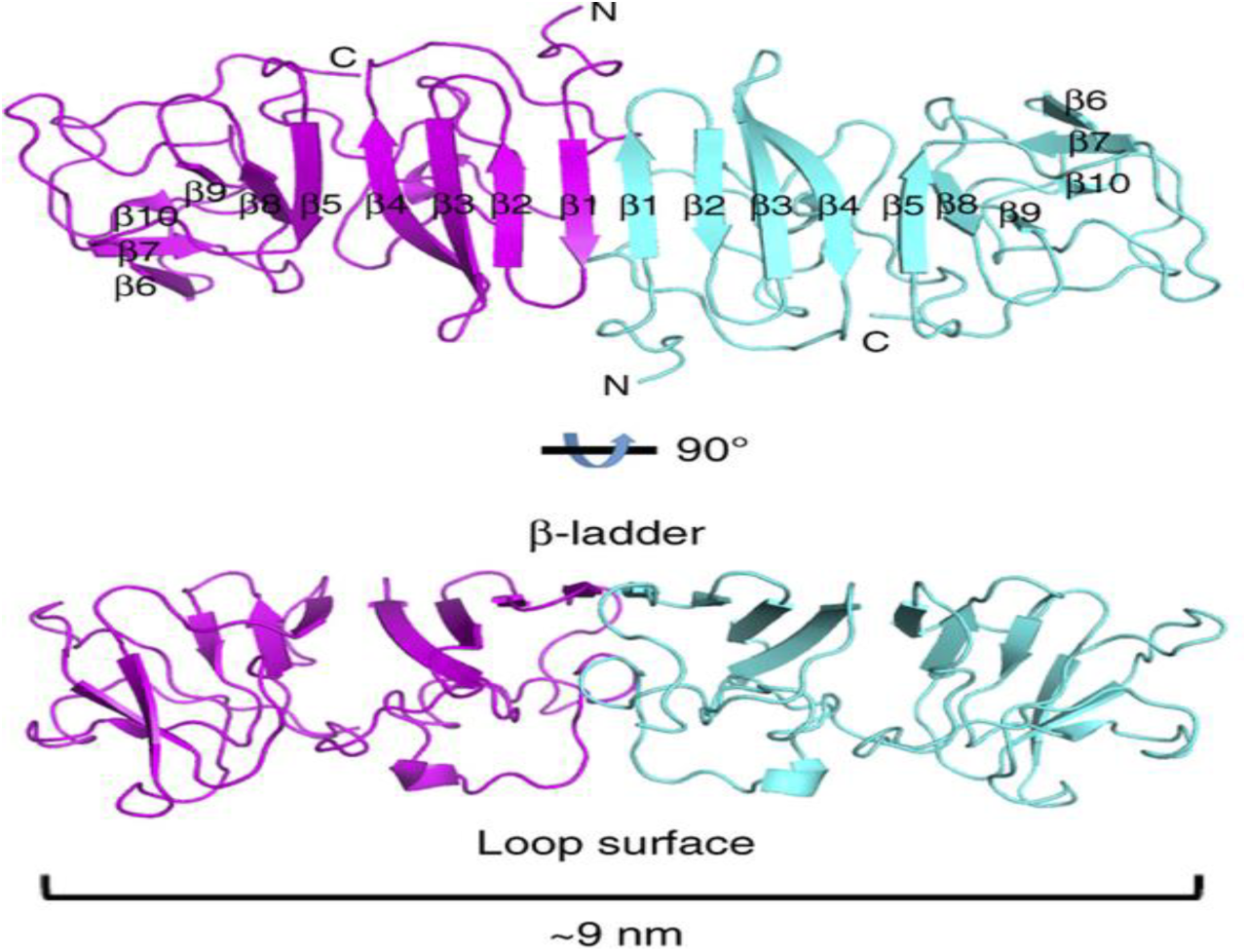
ZIKV NS1 Structure [2]

Most of those inter-strand loops are short, except for a long 'spaghetti loop' between β4 and β5 that lacks secondary structure. In the below figure, a potential N-linked glycosylation site that is highly conserved in the Flaviviridae family is located in the β3–β4 loop [2]. Glycosylation sites are indicated with green hexagons, and disulphide bonds are indicated with yellow circles.

**Figure 6.**
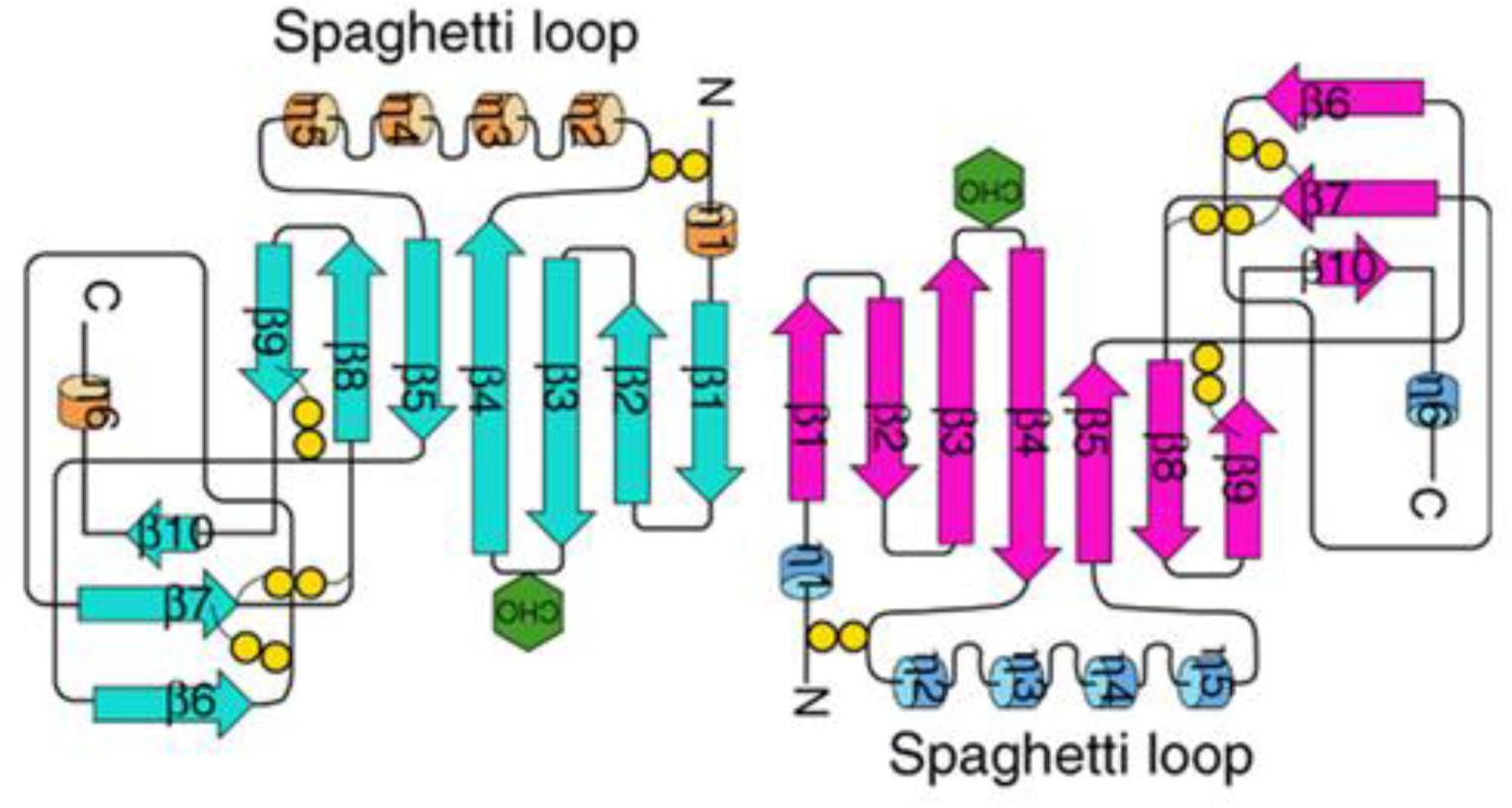
ZIKV NS1 topology diagram [2]

## II. Proposed Method And Implementation

### 2.1 Overview of UCSF Chimera

The Chimera was, according to Greek mythology, a monstrous fire-breathing hybrid creature of Lycia in Asia Minor, composed of the parts of more than one animal. It is usually depicted as a lion, with the head of a goat arising from its back, and a tail that might end with a snake's head.

The term chimera has come to describe any mythical or fictional animal with parts taken from various animals, or to describe anything composed of very disparate parts, or perceived as wildly imaginative, implausible, or dazzling.

In our context and aptly inspired by the word's origin, Chimera is an open-source extensible molecular modeling system developed by University of California, San Francisco (UCSF). Chimera’s primary programming language is Python.

Chimera is divided into a core and extensions. The core provides basic services and molecular graphics capabilities. All higher level functionality is provided through extensions. Extensions can be integrated into the Chimera menu system, and can present a separate graphical user interface as needed using the Tkinter, Tix, and/or Pmw toolkits [4].

The Chimera core consists of a C** layer that handles time-critical operations (e.g., graphics rendering) and a Python layer that handles all other functions. All significant C** data and functions are made accessible to the Python layer. Core capabilities include molecular file input/output, molecular surface generation using the Michel Sanner's Molecular Surface (MSMS) algorithm, and aspects of graphical display such as wire-frame, ball-and-stick, ribbon, and sphere representations, transparency control, near and far clipping planes, and lenses. Another core service is maintenance and display of the current selection. Extensions can query for the contents of the selection [4].

Extensions are written either entirely in Python or in a combination of Python and C/C ++ (the latter using a shared library loaded at runtime). Extensions can be placed in the Chimera installation directory (which would make the extension available to all users) or in the user’s own file area. Extensions are loaded on demand, typically when the user accesses a menu entry that starts the extension [4].

### 2.2. Need for 3D Ramachandran Plot

To help visualize the features of high-fidelity Ramachandran plots, we have found it helpful to look beyond the common two-dimensional ϕ, ψ-plot, which for a large dataset does not serve very well to convey well the true nature of the distribution. In particular, when a large subset of the observations is found very narrowly distributed within one small region, (such as occurs in the α-helical region) this is not well seen in the simple plot because the data points occlude one another.

Three-dimensional versions of the Ramachandran plot, with the third dimension representing observations [1], when coloured and scaled, gives a much more compelling impression of the proportions of residues in the different parts of the Ramachandran plot. It also helps us in visualizing the maximum density zone among all the local maxima.

### 2.3. Working Principle

Figure 9 illustrates the basic working principle of generating a two dimensional Ramachandran plot by running python scripts in UCSF Chimera,

**Figure 7.**
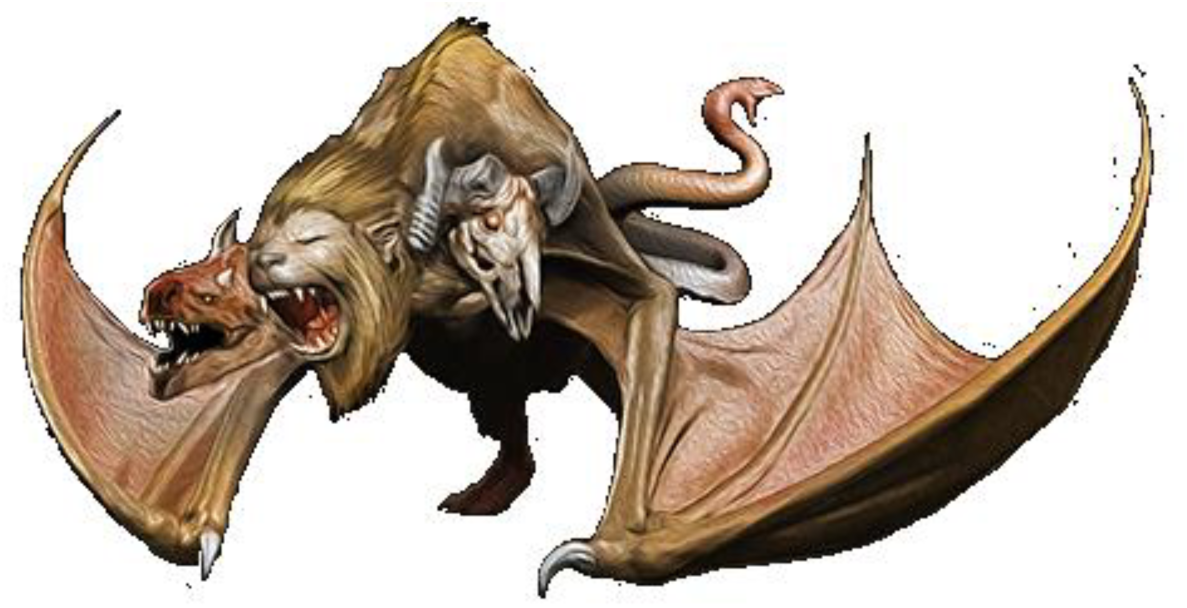
Mythological Chimera

**Figure 8.**
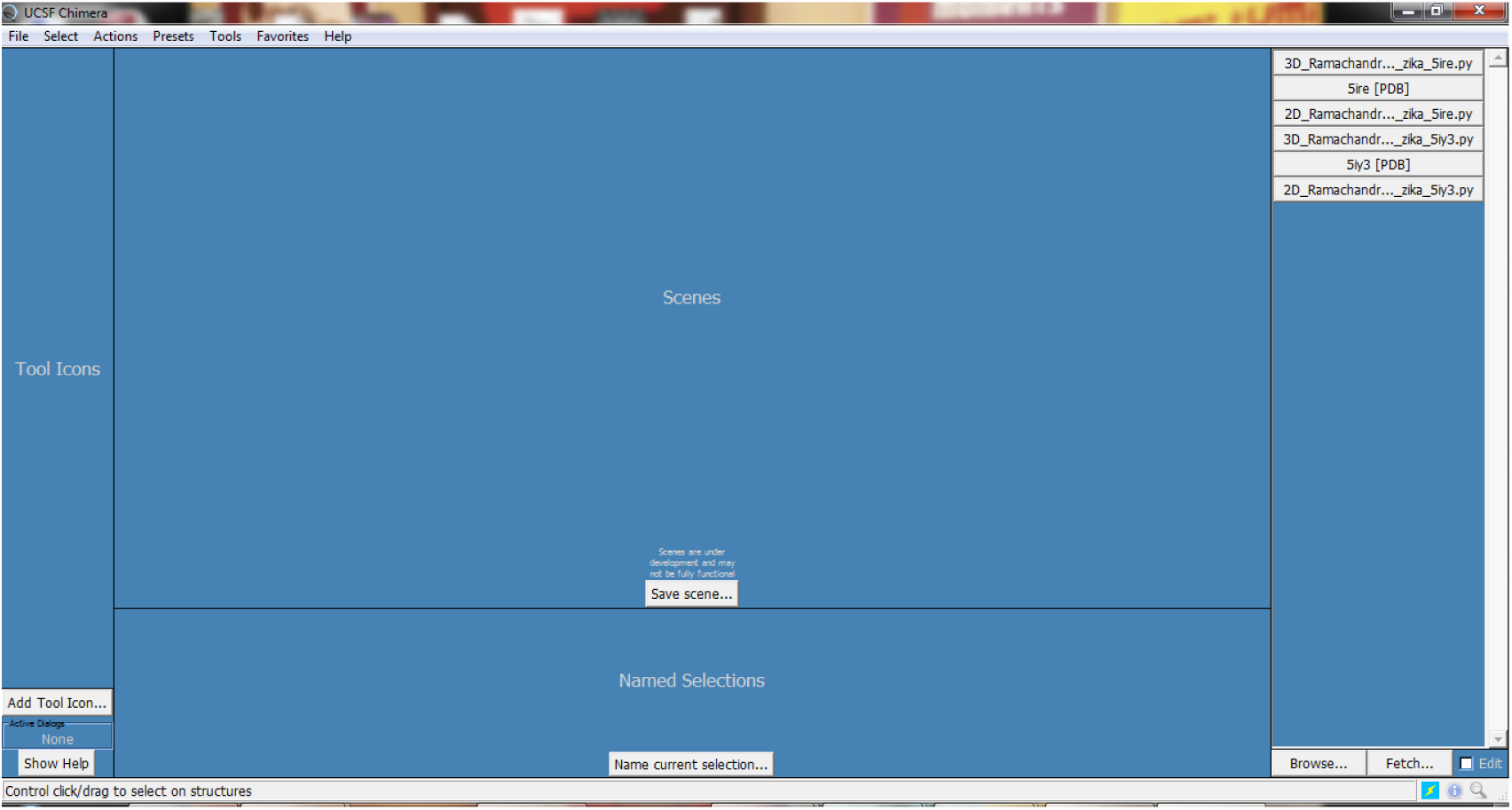
Desktop panel of UCSF Chimera

**Figure 9.**
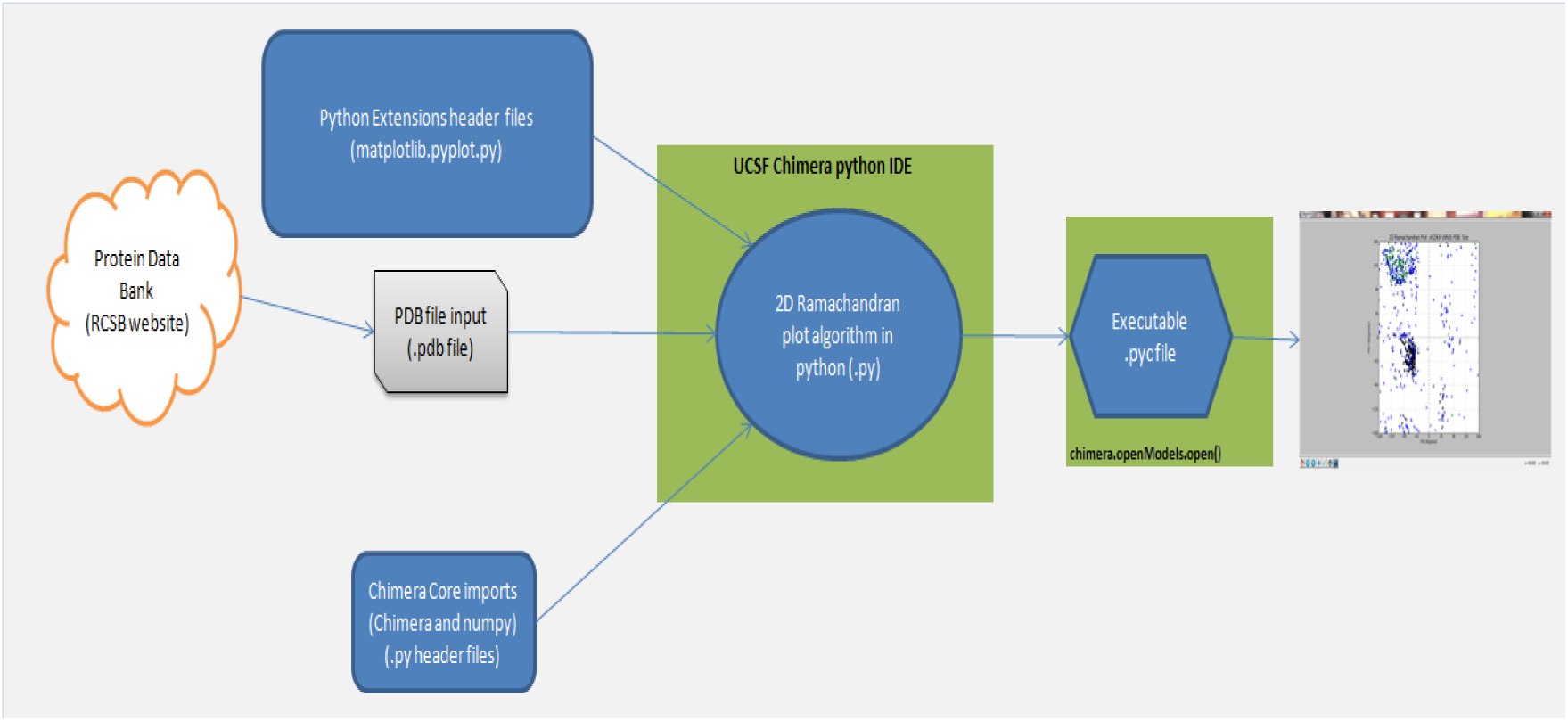
Flow Diagram of 2D Ramachandran plot

Similar flow logic has been used to generate 3D Ramachandran Plot as shown below,

**Figure 10.**
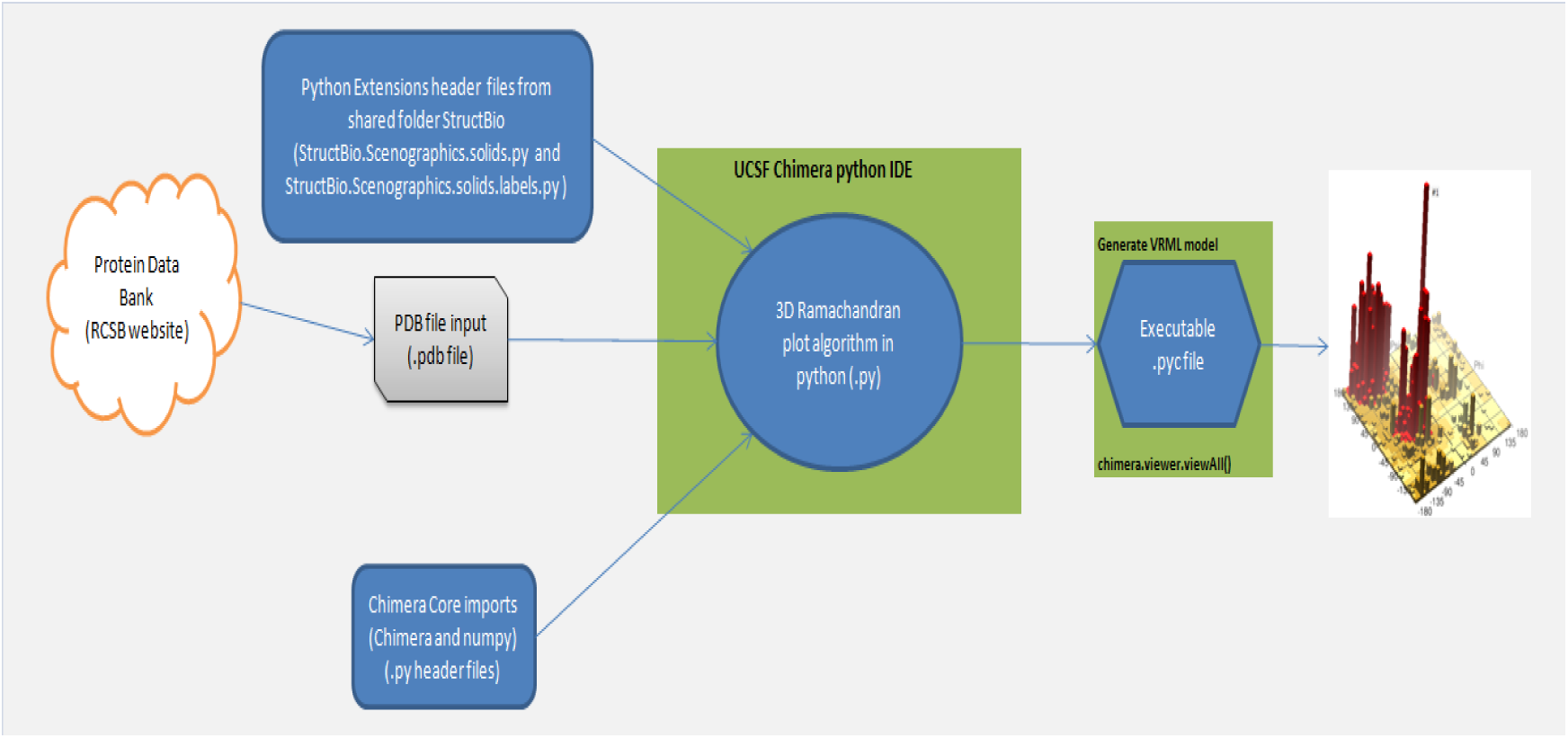
Flow Diagram of 3D Ramachandran plot

The flow diagrams illustrate the following steps:

Step1: Extract the PDB files from RCSB website. For Zika virus, we have two PDB files namely 5IRE.pdb and 5IY3.pdb
Step2: Copy the .pdb file and .py files to the base directory of Chimera tool in Program files.
Step3: Open front panel of Chimera tool and fetch the .py files from the base directory for automated execution.
Step4: Visualize the result output as 2D or 3D plot as per file.

#### 2.3.1 Algorithm for 2D Ramachandra Plot

**Input:** 5ire.pdb/5iy3.pdb

**Output:** 2D Ramachandran plot of 5ire/5iy3

**Steps:**

1. Import Core header files from Chimera and Numpy folders
2. Import Extensive header files pyplot from matplotlib folder
3. Define a function ramaPlot with pdb id as input parameter and image.png file save path as output
4. Set subplot,title,label,limits,ticks,grid,lines
5. Initialize color,shape and lists

//List indicies: 0 for Helix, 1 for Strand, and 2 for Coil
//Marker colors k,g,b for Helix, Strand, and Coil respectively
//Marker shapes o,s,d for Helix, Strand, and Coil respectively
6. Generate the phi, psi coordinate lists (Skip if residue has no phi, psi attributes)
7. Drop in the markers
8. Save file if the saveFilePath parameter is not the empty string
9. Stop ramaPlot definition
10. Call chimera.openModels.open(“5ire", type="PDB”)[0] to get the file from base directory //5iy3
11. Pass it to call function ramaPlot(prot.residues, “5ire”) //5iy3
12. Display the plot in 2D.

#### 2.3.2 Algorithm for 3D Ramachandra Plot

**Input:** 5ire.pdb/5iy3.pdb

**Output:** 3D Ramachandran plot of 5ire

**Steps:**

1. Import Core header files from Chimera and Numpy folders
2. Import Extensive header files solids and labels from StructBio folder
3. Define maxHeight to 64. (maxHeight does not affect the plot unless the largest bucket value is higher than maxHeight in which case the output is scaled so that the “tallest spire" is only maxHeight units high.)
4. Return VRML string that draws a set of boxes (36 * 36). Each box serves as the color coded bucket.
5. Define baseColor and baseHeight
6. For ii in range of 36 Put in cubes making up the base for each jj in range of 36

Put in boxes for bucket values for jj in range of 36
Generate the histogram by adding box for each (i,j) depending on the frequency
Change color code as per the boxed histogram height.
7. Put in the axes in a separate model
8. Take the .pdb file name input and fetch it from base directory using chimera.openModels.open(“5ire", type="PDB”) //5iy3
9. Look at all residues of the protein rather than a single side chain
10. Display the plot in 3d using VRML viewer an rotate manually to get the desired viewing angle.

### 2.4. Graphical Results

Following 3D plot and 2D plot has been taken after execution of the python code in Chimera.

#### 2.4.1 Plot 1: 3D Ramachandran plot for cryo-EM Zika Virus

**Figure 11.**
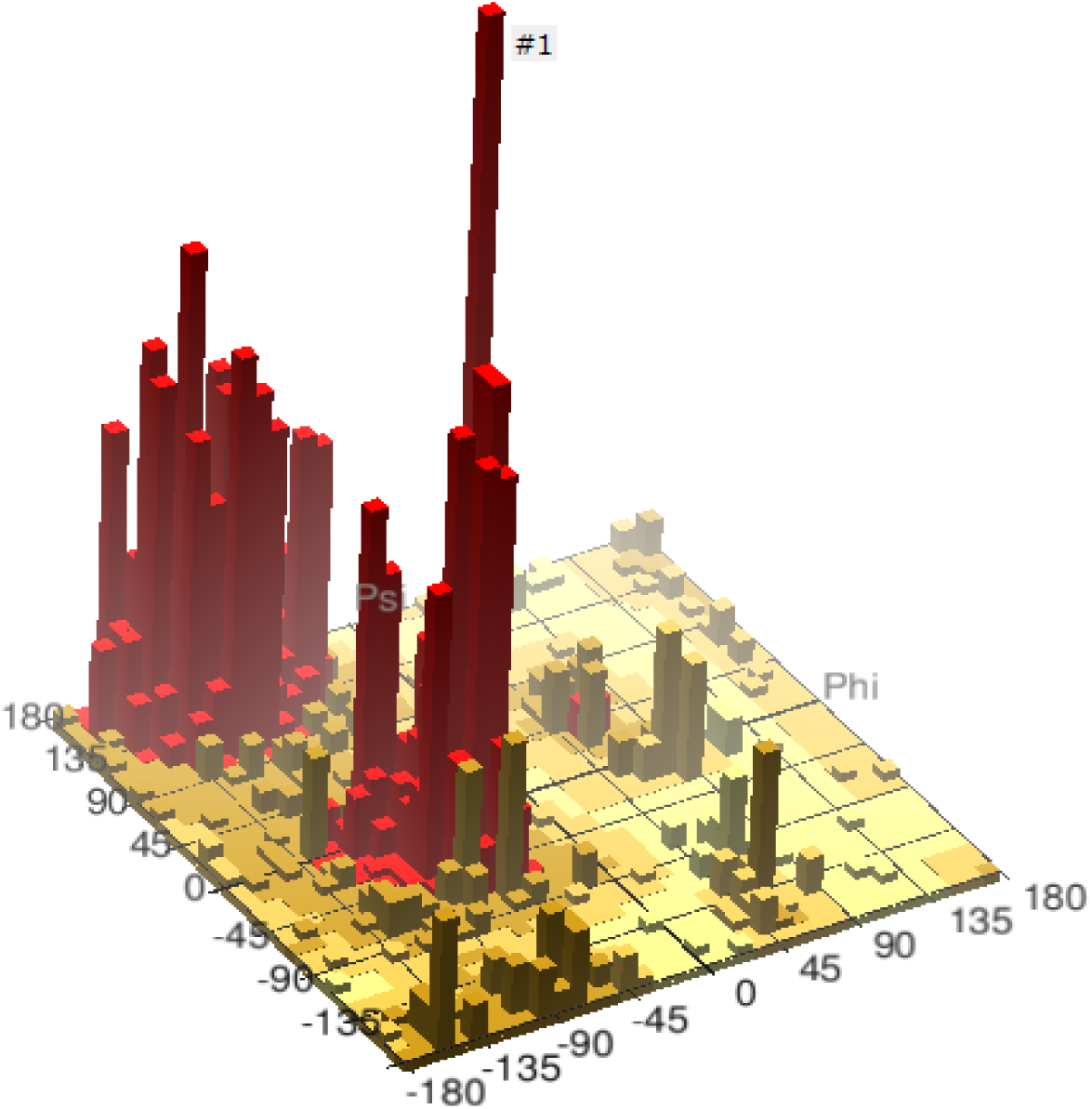
Representing 3D Ramachandran plot 5IRE.pdb

#### 2.4.2 Plot 2: 3D Ramachandran plot for Zika Virus NS1 Protein

**Figure 12.**
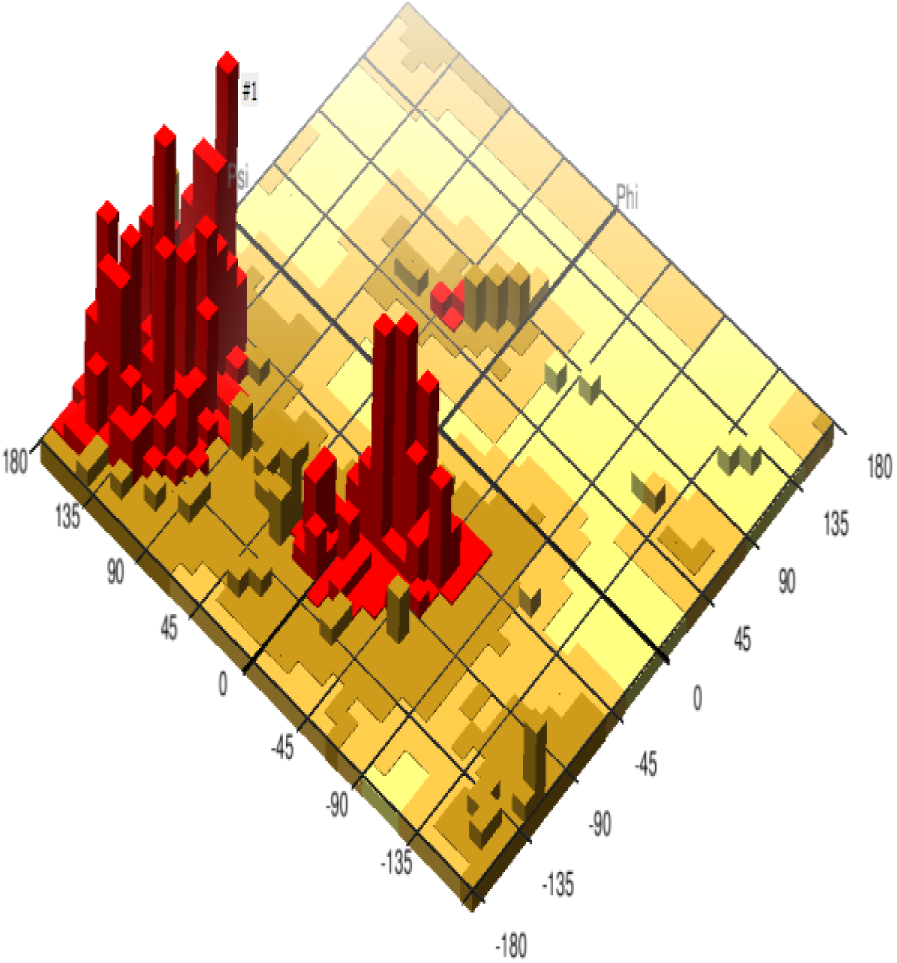
Representing 3D Ramachandran plot 5IY3.pdb

#### 2.4.3 Plot 3: 2D Ramachandran plot for cryo-EM Zika Virus

**Figure 13.**
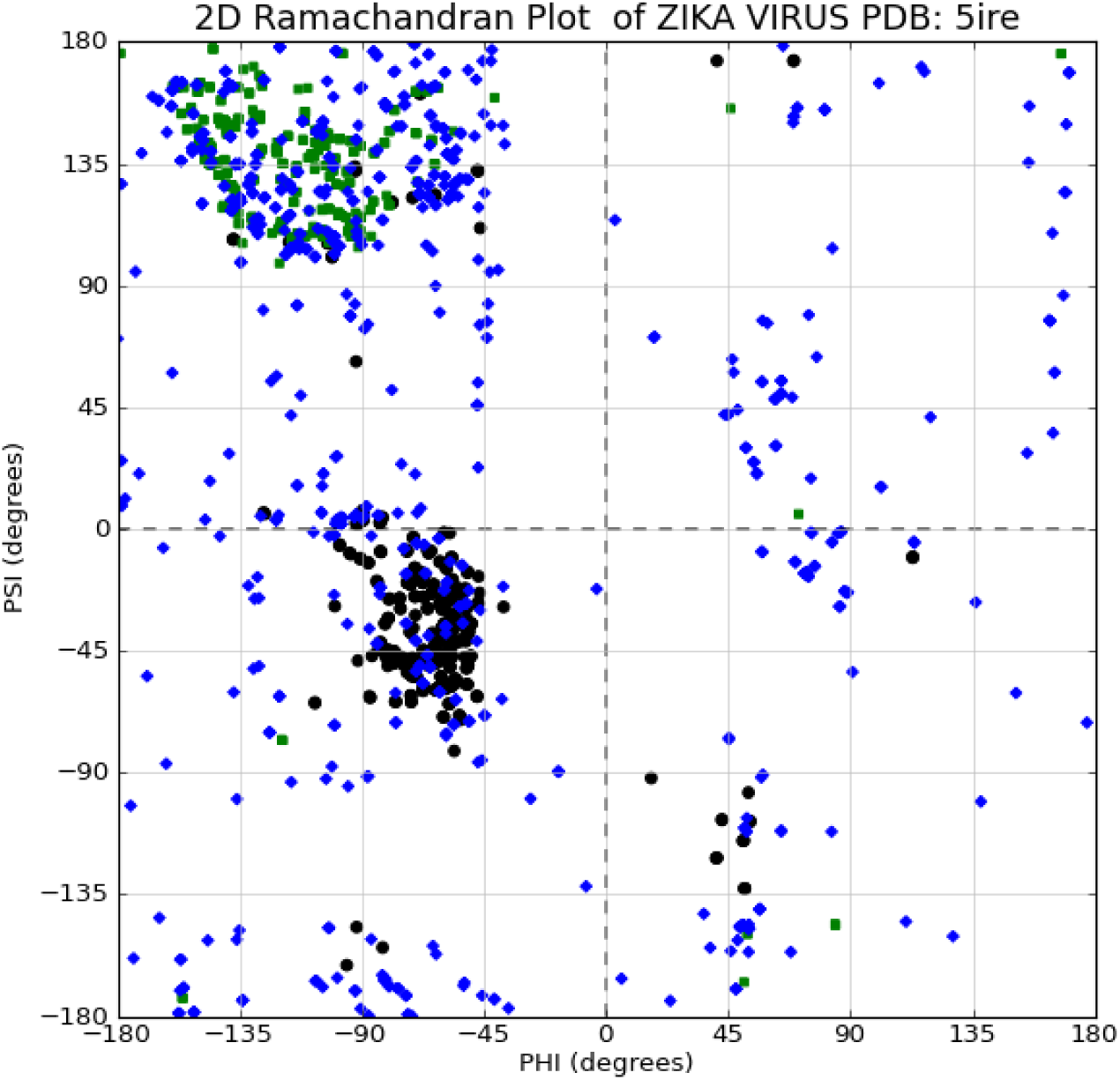
Representing 2D Ramachandran plot 5IRE.pdb

#### 2.4.4 Plot 4: 2D Ramachandran plot for Zika Virus NS1 Protein

**Figure 14.**
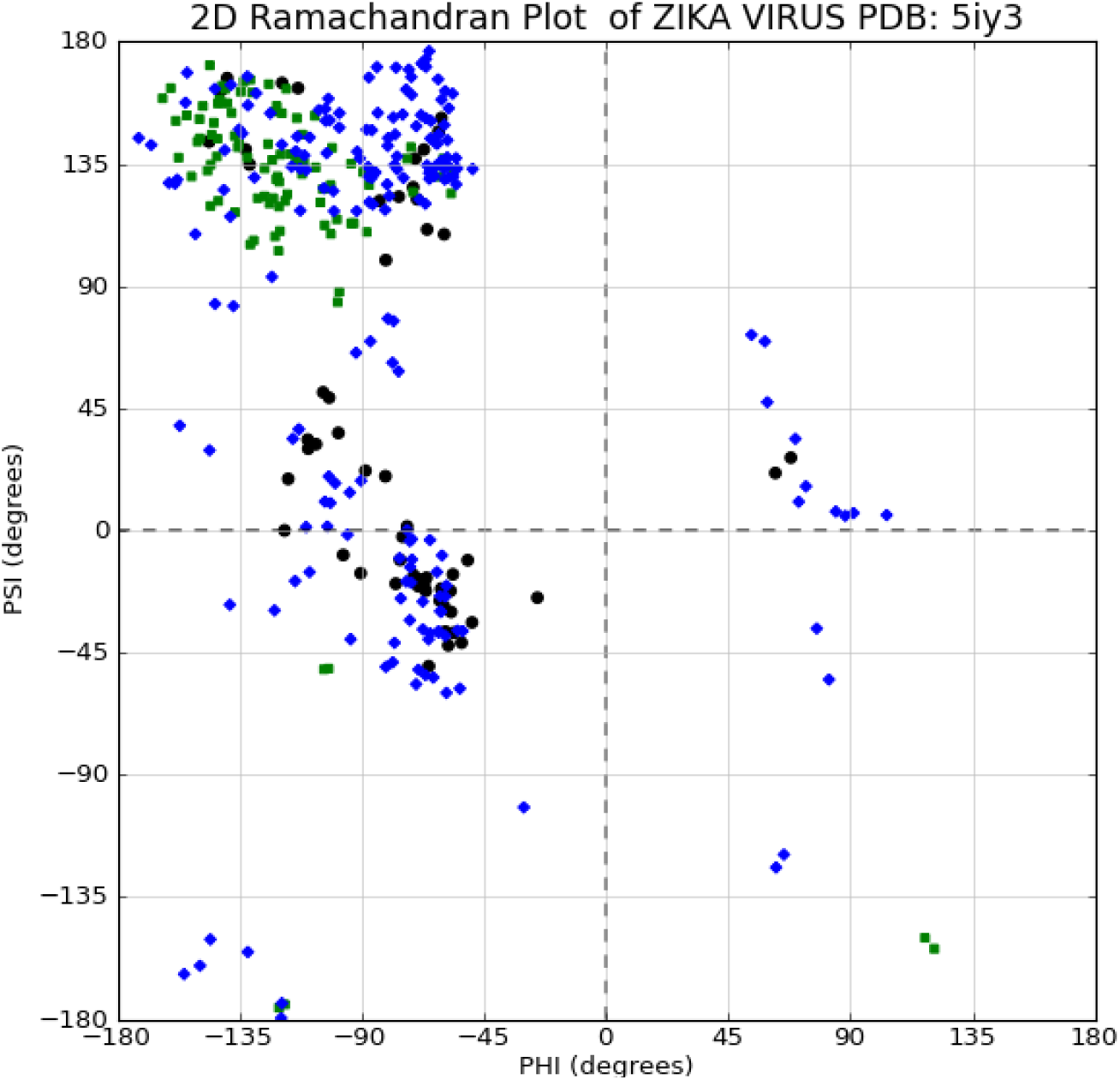
Representing 2D Ramachandran plot 5IY3.pdb

## III. Conclusions

Using our high-fidelity dataset of Zika Virus, the 3D-plot allows several interesting observations. Most obvious, is the titanic and sharp peak resulting from residues in α-helices. This very sharp peak towers over every other portion of the Ramachandran plot, including the other portions of the classically defined alpha-region. This salient observation suggests that the classically defined alpha-region does not behave as a unit.

Moreover we can easily visualize the maximum density zone in the 3D plot marked with #1 rather than large set of scatter point cluster in the 2D graph. The colour codes being graded gives us a clear picture of how the data in protein data bank files are packed over a narrow region.

## IV. Future/Proposed Work

Future works comprise of providing a generalized algorithm to generate a high fidelity 3D plot. This should take box size and maximum height parameters as well. We can even extend this representation as an extension to Protein Data Bank website as a call from JavaScript. The algorithm should also take PDB ID as user inputs rather than hard coding the value in the python script.

Regarding Zika Virus 3D plots, future work should comprise of comparing it with other strains of flavivirus 3D plots to determine the binding positions of the virus and find ways to stop it with suitable antibody.

## V. Acknowledgements

The authors would like to thank Department of Information Technology, Jadavpur University, Salt lake Campus for providing necessary access to scientific literature while conducting this research. The authors acknowledge their gratitude towards Forbes J. Burkowski on his exhaustive literature of structural bioinformatics using UCSF Chimera and Python language[1] which served as the main inspiration for this research.

## Author

**Mayukh Mukhopadhyay** is an ETL developer with over 6 years’ experience in Oracle Data Warehousing. Presently he is working for British Telecommunications Retail Team in TSO platform and pursuing Jadavpur University-TCS collaborative Masters of Engineering in Software Engineering.

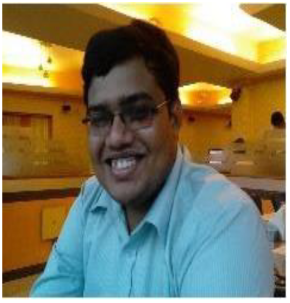

**Dr Parama Bhaumik** is an Assistant Professor in Department of Information Technology, Jadavpur University.

With more than 12 years of academic experience, her areas of interest are Mobile Opportunistic Networks, Bio Inspired Algorithms and Data mining.

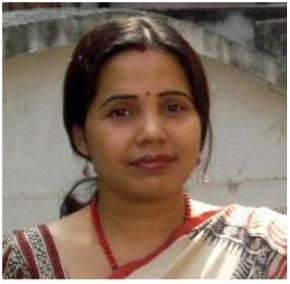

## References

[1] Burkowski, F. J., Computational and Visualization Techniques for Structural Bioinformatics Using Chimera. Chapman & Hall/CRC, 2014.

[2] Zika virus NS1 structure reveals diversity of electrostatic surfaces among flaviviruses, 10.1038/nsmb.3213

[3] The 3.8 Å resolution cryo-EM structure of Zika virus. Science. 2016 Apr 22;352(6284):467–70. doi: 10.1126/science.aaf5316. Epub 2016 Mar 31.

[4] UCSF Chimera--a visualization system for exploratory research and analysis. Pettersen EF, Goddard TD, Huang CC, Couch GS, Greenblatt DM, Meng EC, Ferrin TE. J Comput Chem. 2004 Oct;25(13):1605–12.

[5] Faye O, Freire C, Iamarino A, Faye O, de Oliveira J, Diallo M, Zanotto P, Sall A. (2014) Molecular Evolution of Zika Virus during Its Emergence in the 20th Century. PLOS Neglected Tropical Diseases

[6] Hayes E. (2009) Zika Virus Outside Africa. CDC Emerging Infectious Diseases

[7] Campos GS, Bandeira AC, Sardi SI. (2015) Zika virus outbreak, Bahia, Brazil. CDC Emerging Infectious Diseases

[8] Chambers T, Hahn C, Galler R, Rice C. (1990) Flavivirus genome organization, expression, and replication. Annual Reviews of Microbiology

[9] http://www.cdc.gov/zika/

[10] https://microbewiki.kenyon.edu/index.php/Zika_virus

[11] http://www.vox.com/2016/2/2/10893526/zika-virus-disease-spread-history-cases

[12] Flaviviridae. Swiss Institute of Bioinformatics

